# Validation and invalidation of SARS-CoV-2 main protease inhibitors using the Flip-GFP and Protease-Glo luciferase assays

**DOI:** 10.1101/2021.08.28.458041

**Authors:** Chunlong Ma, Haozhou Tan, Juliana Choza, Yuying Wang, Jun Wang

## Abstract

SARS-CoV-2 main protease (M^pro^) is one of the most extensive exploited drug targets for COVID-19. Structurally disparate compounds have been reported as M^pro^ inhibitors, raising the question of their target specificity. To elucidate the target specificity and the cellular target engagement of the claimed M^pro^ inhibitors, we systematically characterize their mechanism of action using the cell-free FRET assay, the thermal shift-binding assay, the cell lysate Protease-Glo luciferase assay, and the cell-based Flip-GFP assay. Collectively, our results have shown that majority of the M^pro^ inhibitors identified from drug repurposing including ebselen, carmofur, disulfiram, and shikonin are promiscuous cysteine inhibitors that are not specific to M^pro^, while chloroquine, oxytetracycline, montelukast, candesartan, and dipyridamole do not inhibit M^pro^ in any of the assays tested. Overall, our study highlights the need of stringent hit validation at the early stage of drug discovery.

**Graphical abstract:** 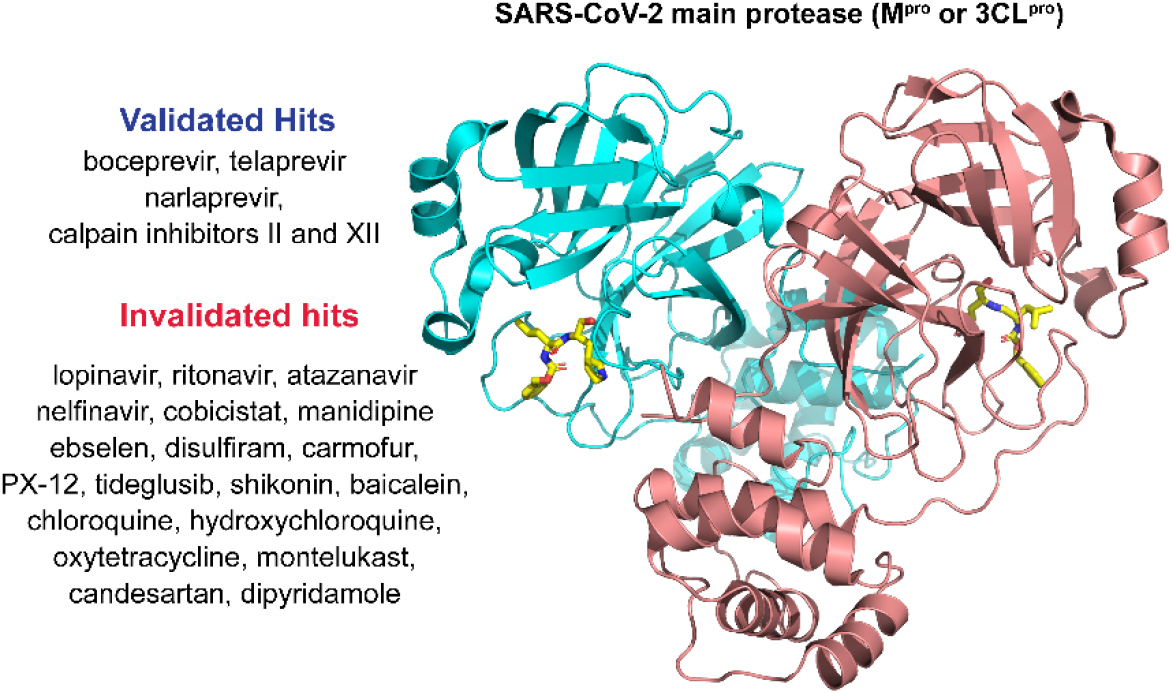

Flip-GFP and Protease-Glo luciferase assays, coupled with the FRET and thermal shift binding assays, were applied to validate the reported SARS-CoV-2 M^pro^ inhibitors.

## 1. INTRODUCTION

SARS-CoV-2 is the causative agent for COVID-19, which infected 221 million people and led to 4.44 million deaths as of August 23, 2021. SARS-CoV-2 is the third coronavirus that causes epidemics and pandemics in human. SARS-CoV-2, along with SARS-CoV and MERS-CoV, belong to the β genera of the coronaviridae family^1^. SARS-CoV-2 encodes two viral cysteine proteases, the main protease (M^pro^) and the papain-like protease (PL^pro^), that mediate the cleavage of viral polyproteins pp1a and pp1ab during viral replication^2, 3^. M^pro^ cleaves at more than 11 sites at the viral polyproteins and has a high substrate preference for glutamine at the P1 site^4^. In addition, the M^pro^ is highly conserved among coronaviruses that infect human including SARS-CoV-2, SARS-CoV, MERS-CoV, HCoV-OC43, HCoV-NL63, HCoV-229E, and HCoV-HKU1. For these reasons, M^pro^ becomes a high-profile drug target for the development of broad-spectrum antivirals. Structurally disparate compounds including FDA-approved drugs and bioactive compounds have been reported as M^pro^ inhibitors^5-7^, several of which also have antiviral activity against SARS-CoV-2^8-10^.

FRET assay is the gold standard assay for protease and is typically used as a primary assay for the screening of M^pro^ inhibitors. However, the FRET assay conditions used by different groups vary significantly in terms of the protein and substrate concentrations, pH, reducing reagent, and detergent. Reducing reagent is typically added in the assay buffer to prevent the non-specific oxidation or alkylation of the catalytic C145 in M^pro^. Nonetheless, many studies do not include reducing reagents in the FRET assay buffer, leading to debatable results^8^. Regardless of the assay condition, FRET assay is a cell free biochemical assay, which does not mimic the cellular environment; therefore, the results cannot be used to accurately predict the cellular activity of the M^pro^ inhibitor or the antiviral activity. Moreover, one limiting factor for M^pro^ inhibitor development is that the cellular activity has to be tested against infectious SARS-CoV-2 in BSL-3 facility, which is inaccessible to many researchers. For these reasons, there is a pressing need of secondary M^pro^ target-specific assays that can closely mimic the cellular environment and be used to rule out false positives.

In this study, we report our findings of validating or invalidating the literature reported M^pro^ inhibitors using the cell lysate Protease-Glo luciferase assay and the cell-based Flip-GFP assay, in conjunction to the cell-free FRET assay and thermal shift-binding assay. The purpose is to elucidate their target specificity and cellular target engagement. The Protease-Glo luciferase assay was developed in this study, and the Flip-GFP assay was recently developed by us and others^11-14^. Our results have collectively shown that majority of the M^pro^ inhibitors identified from drug repurposing screening including ebselen, carmofur, disulfiram, and shikonin are promiscuous cysteine inhibitors that are not specific to M^pro^, while chloroquine, oxytetracycline, montelukast, candesartan, and dipyridamole do not inhibit M^pro^ in any of the assays tested. The results presented herein highlight the pressing need of stringent hit validation at the early stage of drug discovery to minimize the catastrophic failure in the following translational development.

## 2. RESULTS AND DISCUSSION

### 2.1. Assay validation using GC-376 and rupintrivir as positive and negative controls

The advantages and disadvantages of the cell lysate Protease-Glo luciferase assay and the cell-based Flip-GFP assay compared to the cell free FRET assay are listed in Table 1. To minimize the bias from a particular assay, we apply all these three functional assays together with the thermal shift-binding assay for the hit validation.

**Table 1.** Advantages and disadvantages of the three functional assays used in this study.

|  | Advantages | Disadvantages |
| --- | --- | --- |
| **FRET assay** | • High-throughput | • Compounds that quench the fluorophore will show up as false positives<br>• Assay interference from fluorescent compounds, detergents, and aggregators.<br>• Cannot be used to predict the cellular antiviral activity<br>• No standard condition among scientific community |
| **Flip-GFP assay** | • Can rule out compounds that are cytotoxic, membrane impermeable, or substrates of drug efflux pump<br>• A close mimetic of virus-infected cell<br>• Can be used to predict the cellular antiviral activity<br>• Reveals cellular target engagement | • The assay takes 48 hrs, thus it cannot be used for cytotoxic compounds<br>• Interference from fluorescent compounds |
| **Protease-Glo luciferase assay** | • High-throughput<br>• Reveals cellular target engagement<br>• Can be used to test cytotoxic compounds | • Cannot be used to predict the cellular antiviral activity |

In the cell-based Flip-GFP assay, the cells were transfected with two plasmids, one expresses the SARS-CoV-2 M^pro^, and another expresses the GFP reporter^15^. The GFP reporter plasmid expresses three proteins including the GFP β10-β11 fragment flanked by the K5/E5 coiled coil, the GFP β1-9 template, and the mCherry (Fig. 1A). mCherry serves as an internal control for the normalization of the expression level or the quantification of compound toxicity. In the assay design, β10 and β11 were conformationally constrained in the parallel position by the heterodimerizing K5/E5 coiled coil with a M^pro^ cleavage sequence (AVLQ↓SGFR). Upon cleavage of the linker by M^pro^, β10 and β11 become antiparallel and can associate with the β1-9 template, resulting in the restoration of the GFP signal. In principle, the ratio of GFP/mCherry fluorescence is proportional to the enzymatic activity of M^pro^. The Flip-GFP M^pro^ assay has been used by several groups to characterize the cellular activity of M^pro^ inhibitors^11, 13, 14^.

**Figure 1.**
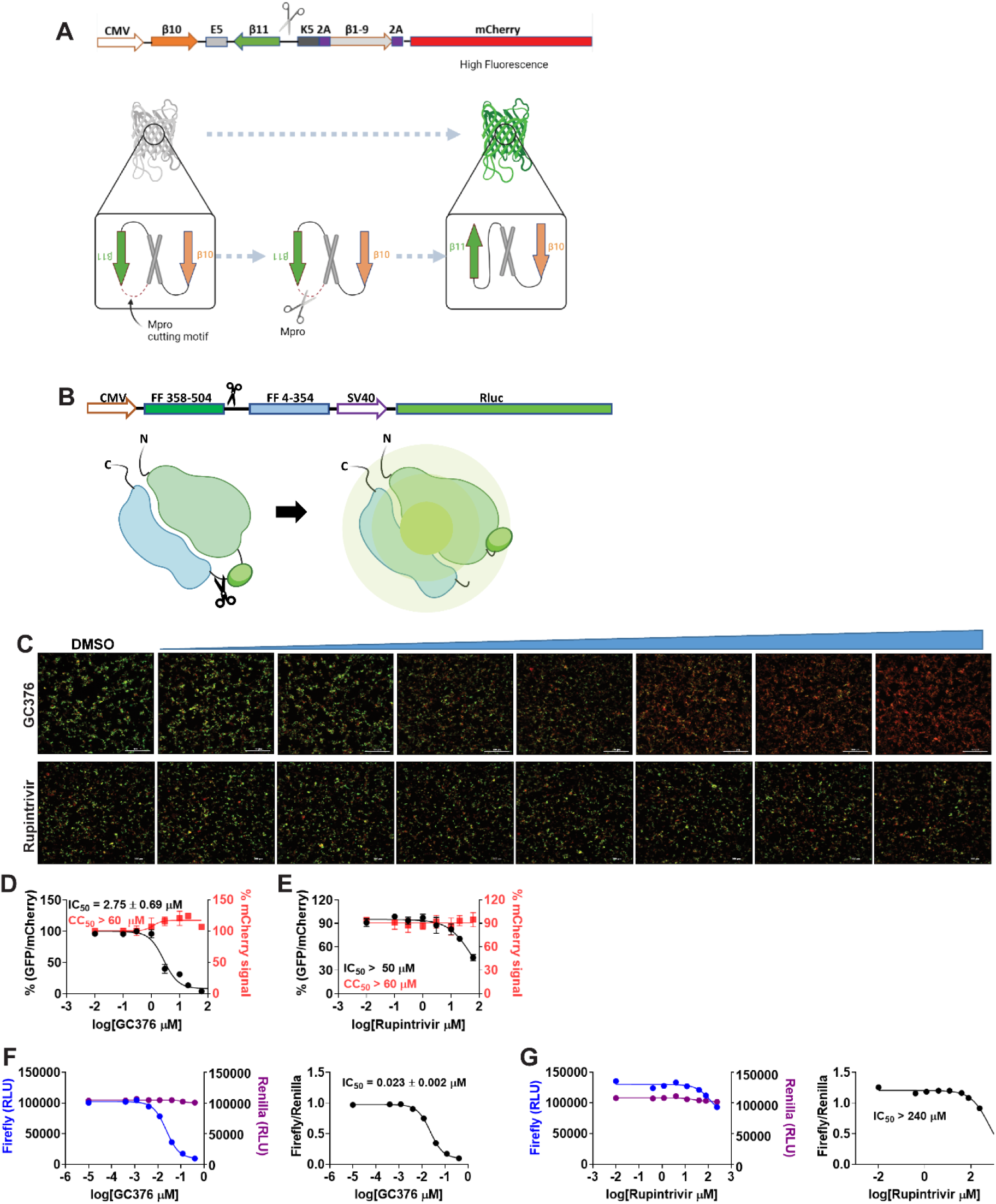
Principles for the Flip-GFP and Protease-Glo luciferase assays and assay validation with control compounds. (A) Assay principle for the Flip-GFP assay. Diagram of the Flip-GFP M^pro^ reporter plasmid is shown. (B) Assay principle for the Protease-Glo luciferase assay. Diagram of pGlo-M^pro^ luciferase reporter in the pGloSensor-30F vector is shown. (C) Representative images from the FlipGFP-M^pro^ assay. Dose-dependent decrease of GFP signal was observed with the increasing concentration of GC-376 (positive control); almost no GFP signal change was observed with the increasing concentration of Rupintrivir (negative control). (D-E) Dose−response curve of the ratio of GFP/mCherry fluorescence with GC-376 and rupintrivir; mCherry signal alone was used to normalize protein expression level or calculate compound cytotoxicity. (F-G) Protease-Glo luciferase assay results of GC-376 and rupintrivir. Left column showed Firefly and Renilla luminescence signals in the presences of increasing concentrations of GC-376 and rupintrivir; Right column showed dose−response curve plots of the ratio of FFluc/Rluc luminescence.

In the cell lysate Protease-Glo luciferase assay, the cells were transfected with pGloSensor-30F luciferase reporter (Fig. 1B)^16^. The pGloSensor-30F luciferase reporter plasmid expresses two proteins, the inactive, circularly permuted firefly luciferase (FFluc) and the active Renilla luciferase (Rluc). Renilla luciferase was included as an internal control to normalize the protein expression level. The firefly luciferase was split into two fragments, the FF 4-354 and FF 358-544. The SARS-CoV-2 M^pro^ substrate cleavage sequence (AVLQ/SGFR) was inserted in between the two fragments. Before protease cleavage, the pGloSensor-30F reporter comprises an inactive circularly permuted firefly luciferase. The cells were lysed at 24 h post transfection, and M^pro^ and the luciferase substrates were added to initiate the reaction. Upon protease cleavage, a conformational change in firefly luciferase leads to drastically increases luminescence. In principle, the ratio of FFluc/Rluc luminescence is proportional to the enzymatic activity of M^pro^.

To calibrate the Flip-GFP and split-luciferase assays, we chose GC-376 and rupintrivir as positive and negative controls, respectively. The IC_50_ values for GC-376 in the Flip-GFP and split-luciferase assays were 2.35 μM and 0.023 μM, respectively (Fig. 1C, D, and F). The IC_50_ value in the Flip-GFP assay is similar to its antiviral activity (Table 2), suggesting the Flip-GFP can be used to predict the cellular antiviral activity. In contrast, rupintrivir showed no activity in either the Flip-GFP (IC_50_ > 50 μM) (Fig. 1C second row and 1E) or the Protease-Glo luciferase assay (IC_50_ > 100 μM) (Fig. 1G), which agrees with the lack of inhibition from the FRET assay (IC_50_ > 20 μM). Nonetheless, rupintrivir was reported to inhibit SARS-CoV-2 replication with an EC_50_ of 1.87 μM using the nanoluciferase SARS-CoV-2 reporter virus (SARS-CoV-2-Nluc) in A549-hACE2 cells^17^ (Table 2). The discrepancy indicates that the mechanism of action of rupintrivir might be independent of M^pro^ inhibition. Overall, the Flip-GFP and Protease-Glo luciferase assays are validated as target-specific assays for SARS-CoV-2 M^pro^.

**Table 2.**
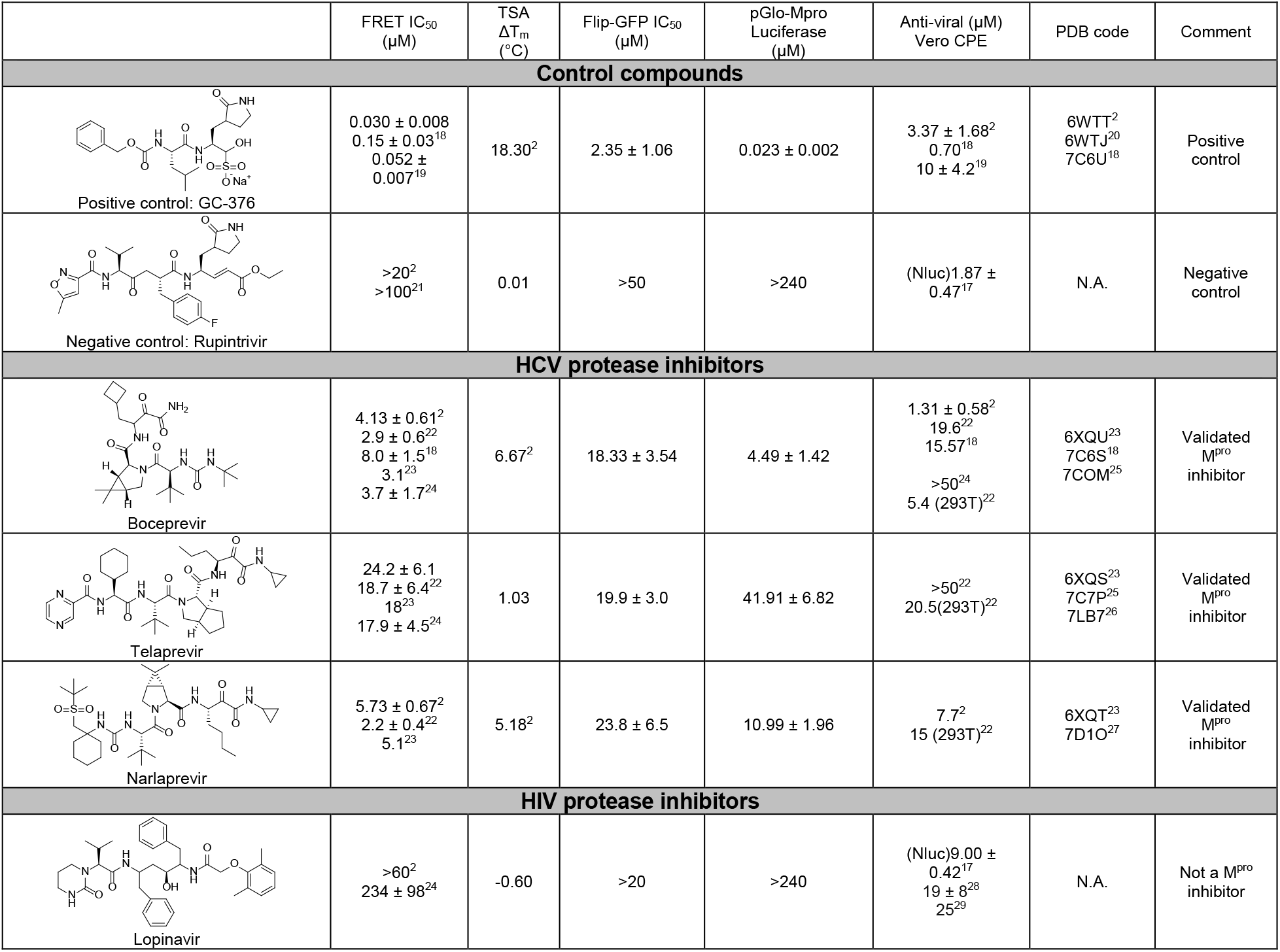
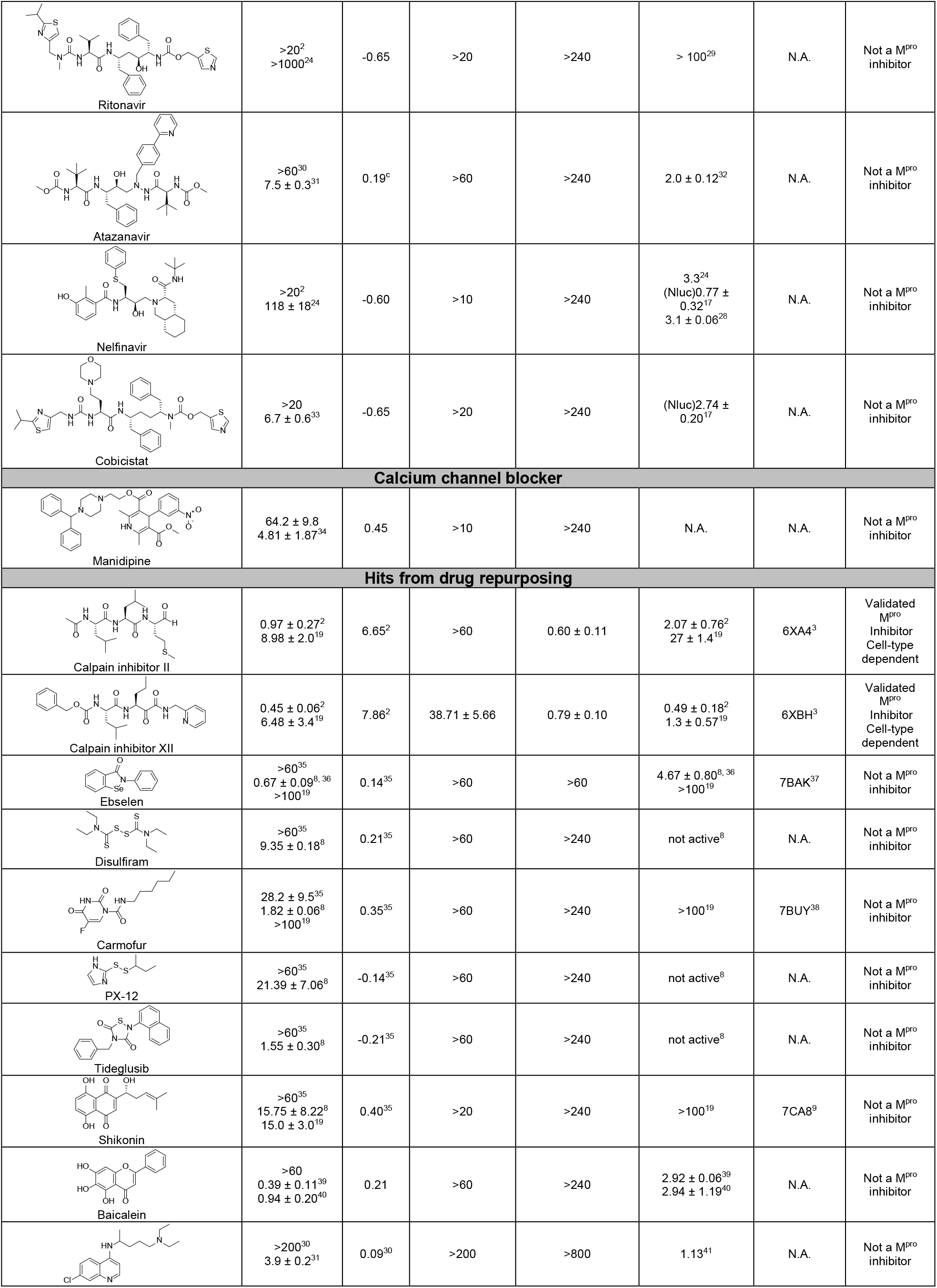
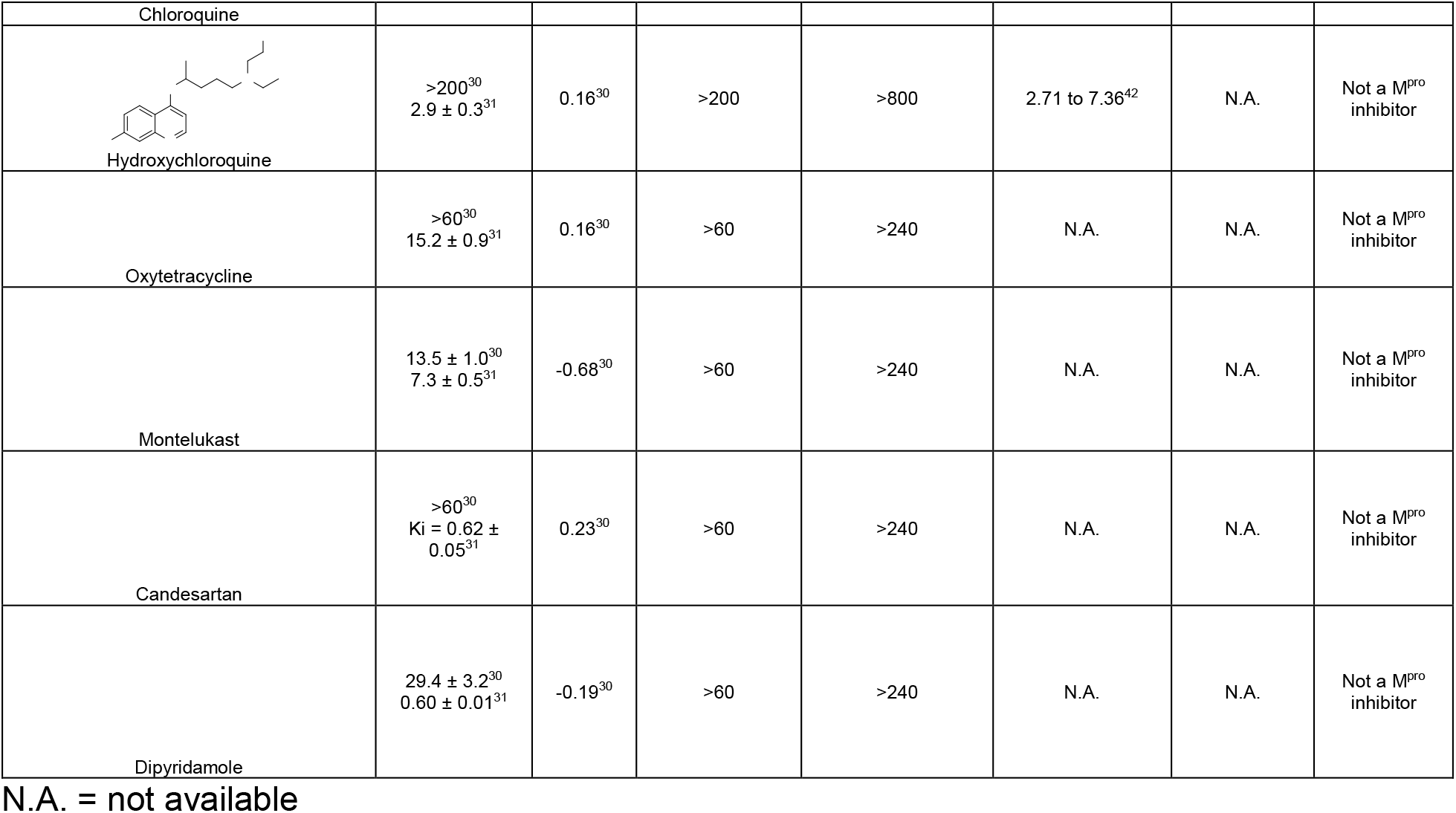
Summary of results.

### 2.2. HCV protease inhibitors

The HCV protease inhibitors have been proven a rich source of SARS-CoV-2 M^pro^ inhibitors^2, 22, 43^. From screening a focused protease library using the FRET assay, we discovered simeprevir, boceprevir, and narlaprevir as SARS-CoV-2 M^pro^ inhibitors with IC_50_ values of 13.74, 4.13, and 5.73 μM, respectively, while telaprevir was less active (31% inhibition at 20 μM)^2^. The binding of boceprevir to M^pro^ was characterized by thermal shift assay and native mass spectrometry. Boceprevir inhibited SARS-CoV-2 viral replication in Vero E6 cells with EC_50_ values of 1.31 and 1.95 μM in the primary CPE and secondary viral yield reduction assays, respectively (Table 2). In parallel, Fu *et al* also reported boceprevir as a SARS-CoV-2 M^pro^ inhibitor with an enzymatic inhibition IC_50_ of 8.0 μM and an antiviral EC_50_ of 15.57 μM^18^. The X-ray crystal structure of M^pro^ with boceprevir was solved, revealing a covalent modification of the C145 thiol by the ketoamide (PDBs: 6XQU^43^, 7C6S^18^, 7COM^25^).

In the current study, we found that boceprevir showed moderate inhibition in the cellular Flip-GFP M^pro^ assay with an IC_50_ of 18.33 μM (Fig. 2A and B), a more than 4-fold increase compared to the IC_50_ in the FRET assay (4.13 μM). The IC_50_ value of boceprevir in the cell lysate Protease-Glo luciferase assay was 4.49 μM (Fig. 2E). In comparison, telaprevir and narlaprevir showed weaker inhibition than boceprevir in both the Flip-GFP and Protease-Glo luciferase assays (Fig. 2A, C, D, F, and G), which is consistent with their weaker potency in the FRET assay (Table 2). Overall, boceprevir, telaprevir, and narlaprevir have been validated as SARS-CoV-2 M^pro^ inhibitors in both the cellular Flip-GFP assay and the cell lysate Protease-Glo luciferase assay. Therefore, the antiviral activity of these three compounds against SARS-CoV-2 are likely due to M^pro^ inhibition. Although the inhibition of M^pro^ by boceprevir is relatively weak compared to GC-376, several highly potent M^pro^ inhibitors were subsequently designed as hybrids of boceprevir and GC-376 including the Pfizer oral drug candidate PF-07321332, which contain the dimethylcyclopropylproline at the P2 substitution^11, 25, 44^.

**Figure 2:**
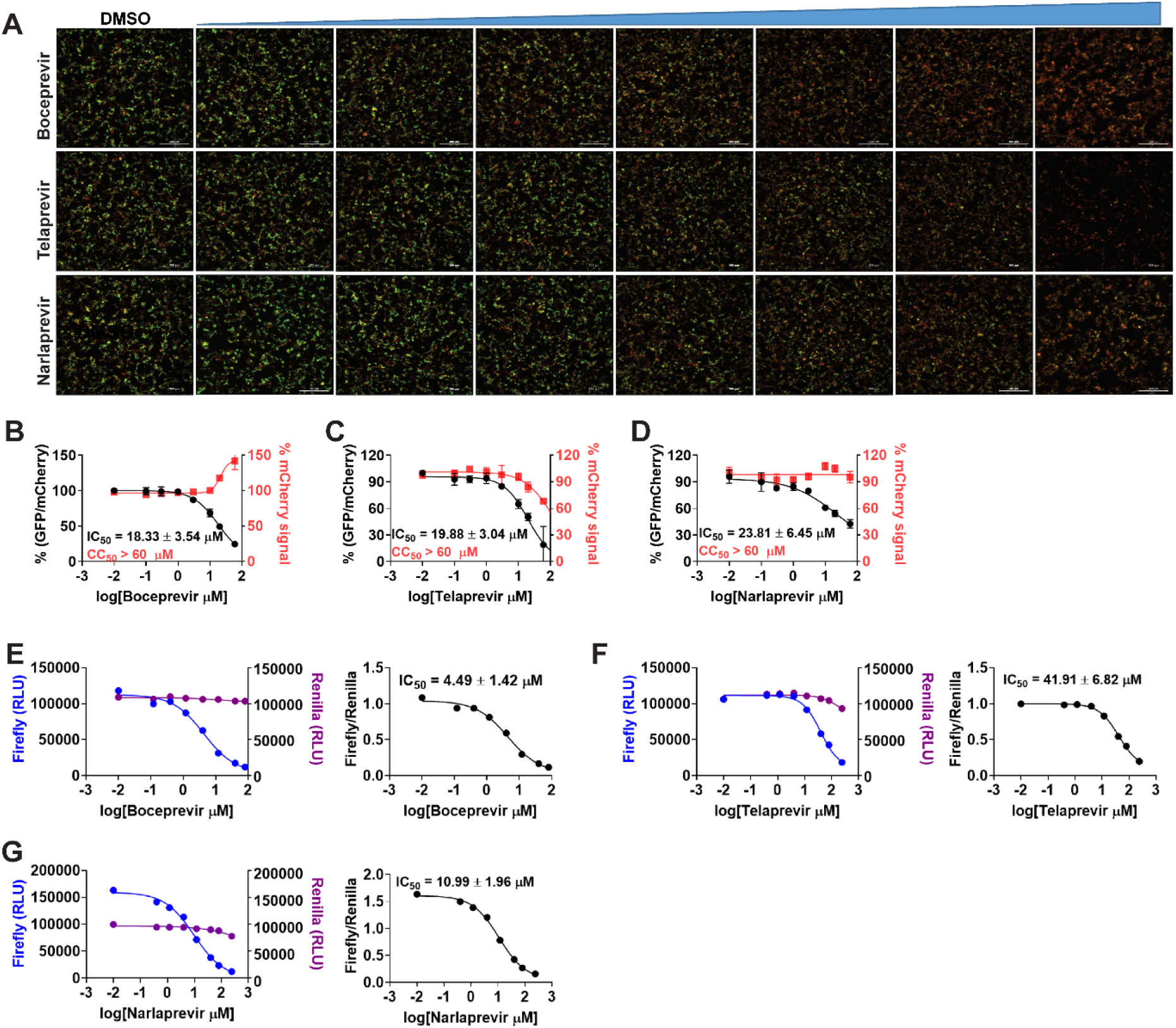
Validation/invalidation of hepatitis C virus NS3/4A protease inhibitors boceprevir, telaprevir, and narlaprevir as SARS CoV-2 M^pro^ inhibitors using the Flip-GFP assay and Protease-Glo luciferase assay. (A) Representative images from the Flip-GFP-M^pro^ assay. Dose-dependent decrease of GFP signal was observed with the increasing concentration of boceprevir, telaprevir or narlaprevir. (B-D) Dose−response curve of the GFP and mCherry fluorescent signals for boceprevir (B), telaprevir (C) or narlaprevir (D); mCherry signal alone was used to normalize protein expression level or calculate compound toxicity. (E-G) Protease-Glo luciferase assay results of boceprevir (E), telaprevir (F) or narlaprevir (G). Left column showed Firefly and Renilla luminescence signals in the presences of increasing concentrations of boceprevir, telaprevir or narlaprevir; Right column showed dose−response curve plots of the ratio of FFluc/Rlu luminescence. Renilla luminescence signal alone was used to normalize protein expression level.

### 2.3. HIV protease inhibitors

HIV protease inhibitors, especially Kaletra, have been hotly pursued as potential COVID-19 treatment at the beginning of the pandemic. Kaletra was first tested in clinical trial during the SARS-CoV outbreak in 2003 and showed somewhat promising results based on the limited data^45^. However, a double-blinded, randomized trial concluded that Kaletra was not effective in treating severe COVID-19^46, 47^. In SARS-CoV-2 infection ferret models, Kaletra showed marginal effect in reducing clinical symptoms, while had no effect on virus titers^48^.

Keletra is a combination of lopinavir and ritonavir. Lopinavir is a HIV protease inhibitor, and ritonavir is used as a booster. Ritonavir does not inhibit the HIV protease and it is a cytochrome P450-3A4 inhibitor^49^. When used in combination, ritonavir can enhance other protease inhibitors by preventing or slowing down the metabolism. In cell culture, lopinavir was reported to inhibit the nanoluciferase SARS-CoV-2 reporter virus with an EC_50_ of 9 μM^17^. In two other studies, lopinavir showed moderate antiviral activity against SARS-CoV-2 activity with EC_50_ values of 19 ± 8 μM^28^ and 25 μM^29^. As such, it was assumed that lopinavir inhibited SARS-CoV-2 through inhibiting the M^pro^. However, lopinavir showed no activity against SARS-CoV-2 M^pro^ in the FRET assay from our previous study (IC_50_ > 60 μM)^2^. Wong et al also showed that lopinavir was a weak inhibitor against SARS-CoV M^pro^ with an IC_50_ of 50 μM^50^. In the current study, we further confirmed the lack of binding of lopinavir to SARS-CoV-2 M^pro^ in the thermal shift assay (ΔT_m_ = - 0.60°C) (Table 2). The result from the Flip-GFP assay was not conclusive as lopinavir was cytotoxic. Lopinavir was not active in the Protease-Glo luciferase assay. Taken together, lopinavir is not a M^pro^ inhibitor.

We also tested additional HIV antivirals including ritonavir, atazanavir, nelfinavir, and cobicistat. Atazanavir and nelfinavir were reported as a potent SARS-CoV-2 antiviral with EC_50_ values of 2.0 ± 0.12^32^ and 0.77 μM^17^ using the infectious SARS-CoV-2 and the nanoluciferase reporter virus (SARS-CoV-2-Nluc), respectively. A drug repurposing screening similar identified nelfinavir as a SARS-CoV-2 antiviral with an IC_50_ of 3.3 μM^24^. Sharma et al showed that cobicistat inhibited M^pro^ with an IC_50_ of 6.7 μM in the FRET assay^33^. Cobicistat was also reported to have antiviral activity against SARS-CoV-2 with an EC_50_ of 2.74 ± 0.20 μM using the SARS-CoV-2-Nluc reporter virus^17^. However, our FRET assay showed that ritonavir, nelfinavir, and cobicistat did not inhibit M^pro^ in the FRET assay (IC_50_ > 20 μM), which was further confirmed by the lack of binding to M^pro^ in the thermal shift assay (Table 2). The results from the Flip-GFP assay were not conclusive due to compound cytotoxicity. None of the compounds showed inhibition in the Protease-Glo luciferase assay.

Collectively, our results have shown that the HIV protease inhibitors including lopinavir, ritonavir, atazanavir, nelfinavir, and cobicistat are not M^pro^ inhibitors. Nonetheless, given the potent antiviral activity of atazanavir and nelfinavir against SARS-CoV-2, it might be interesting to conduct resistance selection to elucidate their drug target(s).

### 2.4. Bioactive compounds from drug repurposing

Several bioactive compounds have been identified as SARS-CoV-2 M^pro^ inhibitors through either virtual screening or FRET-based HTS. We are interested in validating these hits using the Flip-GFP and the Protease-Glo luciferase assays.

Manidipine was identified as a SARS-CoV-2 M^pro^ inhibitor from a virtual screening and was subsequently shown to inhibit M^pro^ with an IC_50_ of 4.81 μM in the FRET assay^34^. No antiviral data was provided. When we repeated the FRET assay, the IC_50_ was 64.2 μM (Table 2). Manidipine also did not show binding to M^pro^ in the thermal shift assay. Furthermore, manidipine showed no activity in either the Flip-GFP assay or the Protease-Glo luciferase assay (Fig. 4A, B, and F). Therefore, our results invalidated manidipine as a SARS-CoV-2 M^pro^ inhibitor.

**Figure 3:**
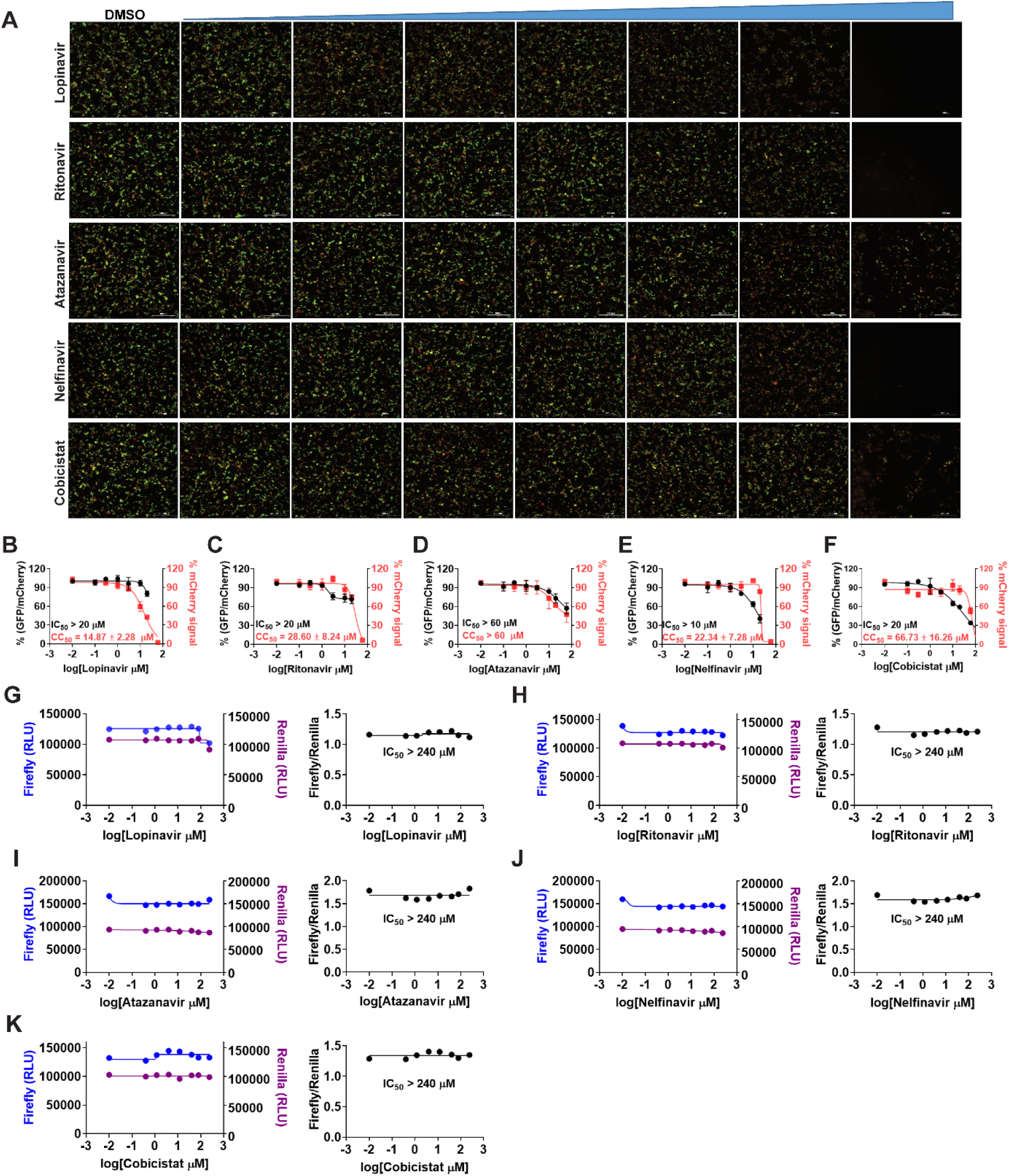
Validation/invalidation of HIV protease inhibitors lopinavir, ritonavir, atazanavir, nelfinavir, and cobicistat as SARS CoV-2 M^pro^ inhibitors using the Flip-GFP assay and Protease-Glo luciferase assay. (A) Representative images from the Flip-GFP-M^pro^ assay. (B-F) Dose−response curve of the GFP and mCherry fluorescent signals for lopinavir (B), ritonavir (C), atazanavir (D), nelfinavir (E), and cobicistat (F); mCherry signal alone was used to normalize protein expression level or calculate compound cytotoxicity. (G-K) Protease-Glo luciferase assay results of lopinavir (G), ritonavir (H), atazanavir (I), nelfinavir (J), and cobicistat (K). Left column showed Firefly and Renilla luminescence signals in the presences of increasing concentrations of lopinavir, ritonavir, atazanavir, nelfinavir, and cobicistat; Right column showed dose−response curve plots of ratio of FFluc/Rluc luminescence. Renilla luminescence signal alone was used to normalize protein expression level. None of the compounds shows significant inhibition in the presence of up to 240 μM compounds.

**Figure 4.**
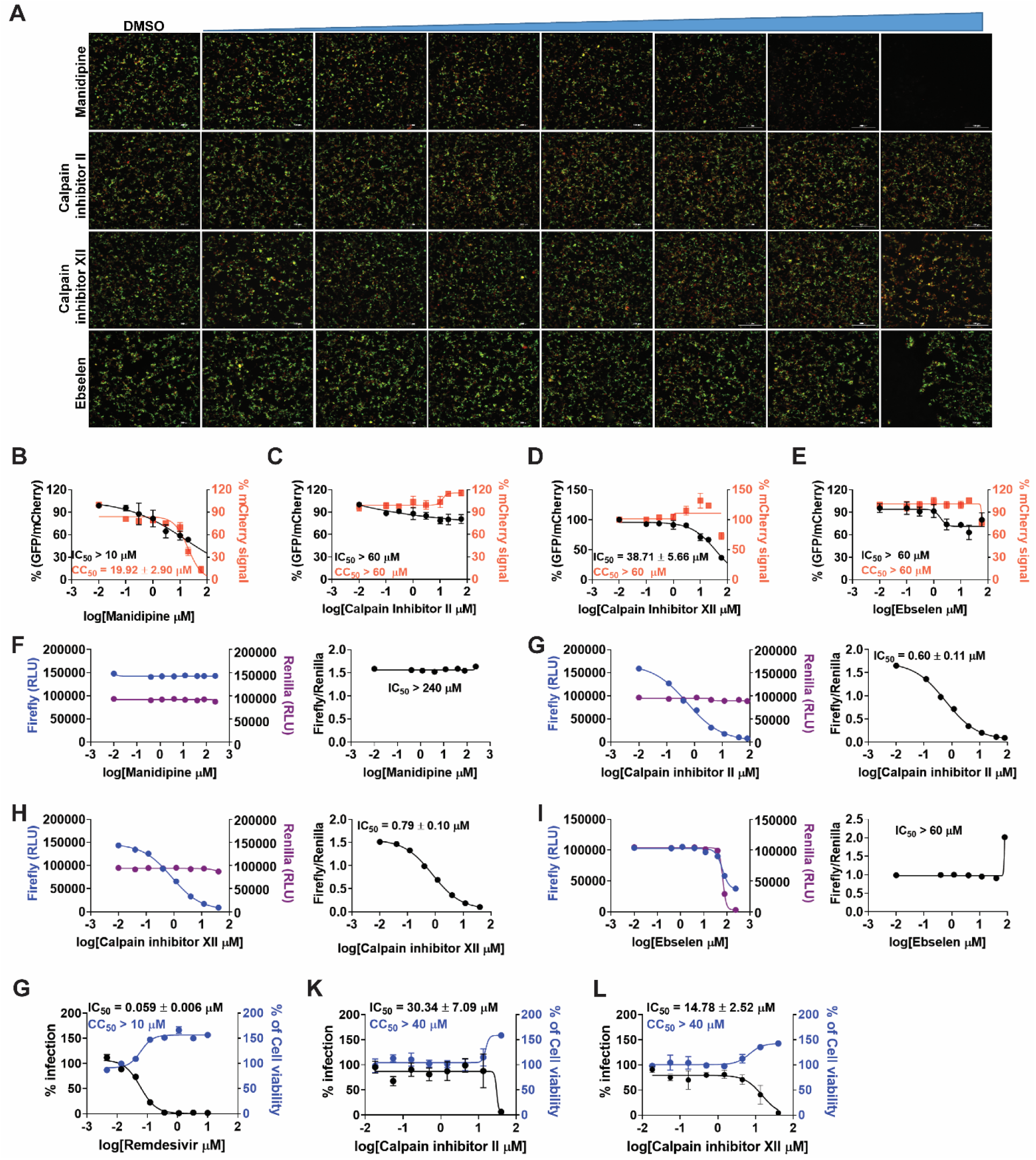
Validation/invalidation of manidipine, calpain inhibitors II and XII, and ebselen as SARS CoV-2 M^pro^ inhibitors using the Flip-GFP assay and Protease-Glo luciferase assay. (A) Representative images from the Flip-GFP-M^pro^ assay. (B-E) Dose−response curve of the GFP and mCherry fluorescent signals for manidipine (B), calpain inhibitor II (C), calpain inhibitor XII (D), and ebselen (E); mCherry signal alone was used to normalize protein expression level or calculate compound cytotoxicity. (F-I) Protease-Glo luciferase assay results of manidipine (F), calpain inhibitor II (G), calpain inhibitor XII (H), and ebselen (I). Left column showed Firefly and Renilla luminescence signals in the presences of increasing concentrations of lopinavir, ritonavir, atazanavir, nelfinavir, and cobicistat; Right column showed dose−response curve plots of the ratio of FFluc/Rluc luminescence. Renilla luminescence signal alone was used to normalize protein expression level. (G-K) Antiviral activity of remdesivir (G), calpain inhibitor II (K), and calpain inhibitor XII (L) against SARS-CoV-2 in Calu-3 cells.

In the same screening which we identified boceprevir as a SARS-CoV-2 M^pro^ inhibitor, calpain inhibitors II and XII were also found to have potent inhibition against M^pro^ with IC_50_ values of 0.97 and 0.45 μM in the FRET assay^2^. Both compounds showed binding to M^pro^ in the thermal shift and native mass spectrometry assays. The Protease-Glo luciferase assay similarly confirmed the potent inhibition of calpain inhibitors II and XII against M^pro^ with IC_50_ values of 0.60 and 0.79 μM, respectively (Fig. 4G, H). However, calpain inhibitor II had no effect on the cellular M^pro^ activity as shown by the lack of inhibition in the Flip-GFP assay (IC_50_ > 60 μM) (Fig. 4A, C), while calpain inhibitor XII showed weak activity (IC_50_ = 38.71 μM) (Fig. 4A, D). A recent study by Liu et al using a M^pro^ trigged cytotoxicity assay similarly found the lack of cellular M^pro^ inhibition by calpain inhibitors II and XII^51^. These results contradict to the potent antiviral activity of both compounds in Vero E6 cells^2^. It is noted that calpain inhibitors II and XII are also potent inhibitors of cathepsin L with IC_50_ values of 0.41 and 1.62 nM, respectively^3^. One possible explanation is that the antiviral activity of calpain inhibitors II and XII against SARS-CoV-2 might be cell type dependent, and the observed inhibition in Vero E6 cells might be due to cathepsin L inhibition instead of M^pro^ inhibition. Vero E6 cells are TMPRSS2 negative, and SARS-CoV-2 enters cell mainly through endocytosis and is susceptible to cathepsin L inhibitors^52^. To further evaluate the antiviral activity of calpain inhibitors II and XII against SARS-CoV-2, we tested them in Calu-3 cells using the immunofluorescence assay (Fig. 4G, K, L). Calu-3 is TMPRSS2 positive and it is a close mimetic of the human primary epithelial cell^53^. As expected, calpain inhibitors II and XII displayed much weaker antiviral activity against SARS-CoV-2 in Calu-3 cells than in Vero E6 cells with EC_50_ values of 30.34 and 14.78 μM, respectively (Fig. 4K, L). These results suggest that the Flip-GFP assay can be used to faithfully predict the antiviral activity of M^pro^ inhibitors. The lower activity of calpain inhibitors II and XII in the Flip-GFP assay and the Calu-3 antiviral assay might due to the competition with host proteases, resulting in the lack of cellular target engagement with M^pro^.

In conclusion, calpain inhibitors II and XII are validated as M^pro^ inhibitors but their antiviral activity against SARS-CoV-2 is cell type dependent. Accordingly, TMPRSS2 positive cell lines such as Calu-3 should be used to test the antiviral activity of calpain inhibitors II and XII analogs.

Ebselen is among one of the most frequently reported promiscuous M^pro^ inhibitors. It was first reported by Yang et al that ebselen inhibits SARS-CoV-2 M^pro^ with an IC_50_ of 0.67 μM and the SARS-CoV-2 replication with an EC_50_ of 4.67 μM^8^. However, it was noted that no reducing reagent was added in the FRET assay, and we reasoned that the observed inhibition might be due to non-specific modification of the catalytic cysteine 145 by ebselen. To test this hypothesis, we repeated the FRET assay with and without reducing reagent DTT or GSH, and found that ebselen completely lost the M^pro^ inhibition in the presence of DTT or GSH^35^. Similarly, ebselen also non-specifically inhibited several other viral cysteine proteases in the absence of DTT including SARS-CoV-2 PL^pro^, EV-D68 2A^pro^ and 3C^pro^, and EV-A71 2A^pro^ and 3C^pro35^. The inhibition was abolished with the addition of DTT. Ebselen also had no antiviral activity against EV-A71 and EV-D68, suggesting that the FRET assay results without reducing reagent cannot be used to predict the antiviral activity. In this study, we found that ebselen showed no inhibition in either the Flip-GFP assay or the split-luciferase assay (Fig. 4A, E, I), providing further evidence for the promiscuous mechanism of action of ebselen. Another independent study by Deval et al using mass spectrometry assay reached similar conclusion that the inhibition of M^pro^ by ebselen is non-specific and inhibition was abolished with the addition of reducing reagent DTT or glutathione ^54^. In contrary to the potent antiviral activity reported by Yang et al, the study from Deval et al found that ebselen was inactive against SARS-CoV-2 in Vero E6 cells (EC_50_ > 100 μM). Lim et al reported that ebselen and disulfiram had synergistic antiviral effect with remdesivir against SARS-CoV-2 in vero E6 cells^55^. It was proposed that ebselen and disulfiram act as zinc ejectors and inhibited not only the PL^pro56^, but also the nsp13 ATPase and nsp14 exoribonuclease activities^55^, further casting doubt on the detailed mechanism of action of ebselen.

Despite the accumulating evidence to support the promiscuous mechanism of action of ebselen, several studies continue to explore ebselen and its analogs as SARS-CoV-2 M^pro^ and PL^pro^ inhibitors^36, 57, 58^. A number of ebselen analogs were designed and found to have comparable enzymatic inhibition and antiviral activity as ebselen. MR6-31-2 had slightly weaker enzymatic inhibition against SARS-CoV-2 M^pro^ compared to ebselen (IC_50_ = 0.824 vs 0.67 μM), however, MR6-31-2 had more potent antiviral activity than ebselen (EC_50_ = 1.78 vs 4.67 μM) against SARS-CoV-2 M^pro^ in Vero E6 cells. X-ray crystallization of SARS-CoV-2 M^pro^ with MR6-31-2 (PDB: 7BAL) and ebselen (PDB: 7BAK) revealed nearly identical complex structures. It was found that selenium coordinates directly to Cys145 and forms a S-Se bond^36^. Accordingly, a mechanism involving hydrolysis of the organoselenium compounds was proposed. Similar to their previous study, the M^pro^ enzymatic reaction buffer (50 mM Tris pH 7.3, 1 mM EDTA) did not include the reducing reagent DTT. Therefore, the M^pro^ inhibition by these ebselen analogs might be non-specific and the antiviral activity might arise from other mechanisms.^36^

Overall, it can be concluded that ebselen is not a specific M^pro^ inhibitor, and its antiviral activity against SARS-CoV-2 might involve other drug targets such as nsp13 or nsp14.

Disulfiram is an FDA-approved drug for alcohol aversion therapy. Disulfiram has a polypharmacology and was reported to inhibit multiple enzymes including urease^59^, methyltransferase^60^, and kinase^59^ through reacting with cysteine residues. Disulfiram was also reported as an allosteric inhibitor of MERS-CoV PL^pro61^. Yang et al reported disulfiram as a M^pro^ inhibitor with an IC_50_ of 9.35 μM. Follow up studies by us and others showed that disulfiram did not inhibit M^pro^ in the presence of DTT. In this study, disulfiram had no inhibition against M^pro^ in either the Flip-GFP assay or the Protease-Glo luciferase assay (Fig. 5A, B, N).

**Figure 5.**
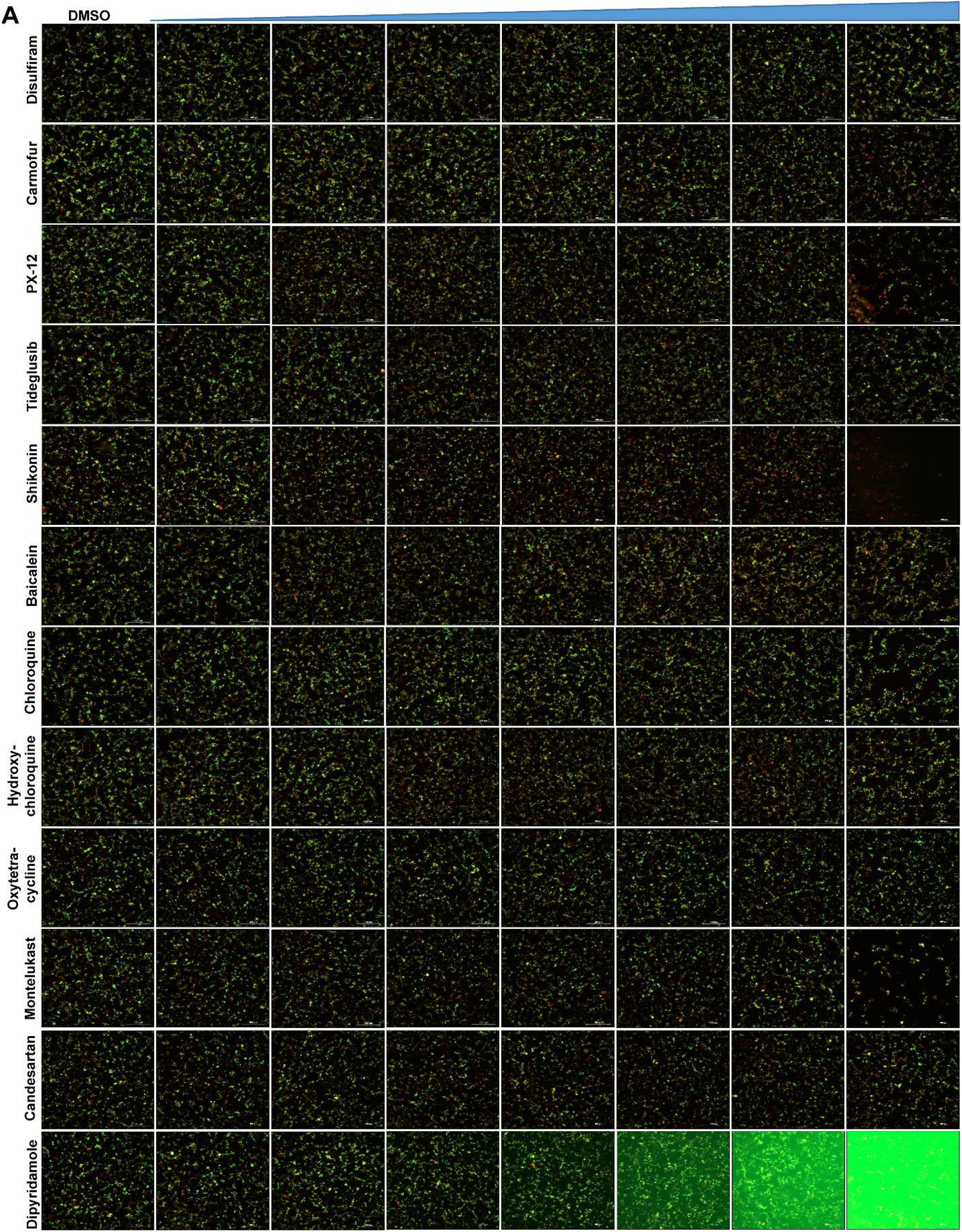
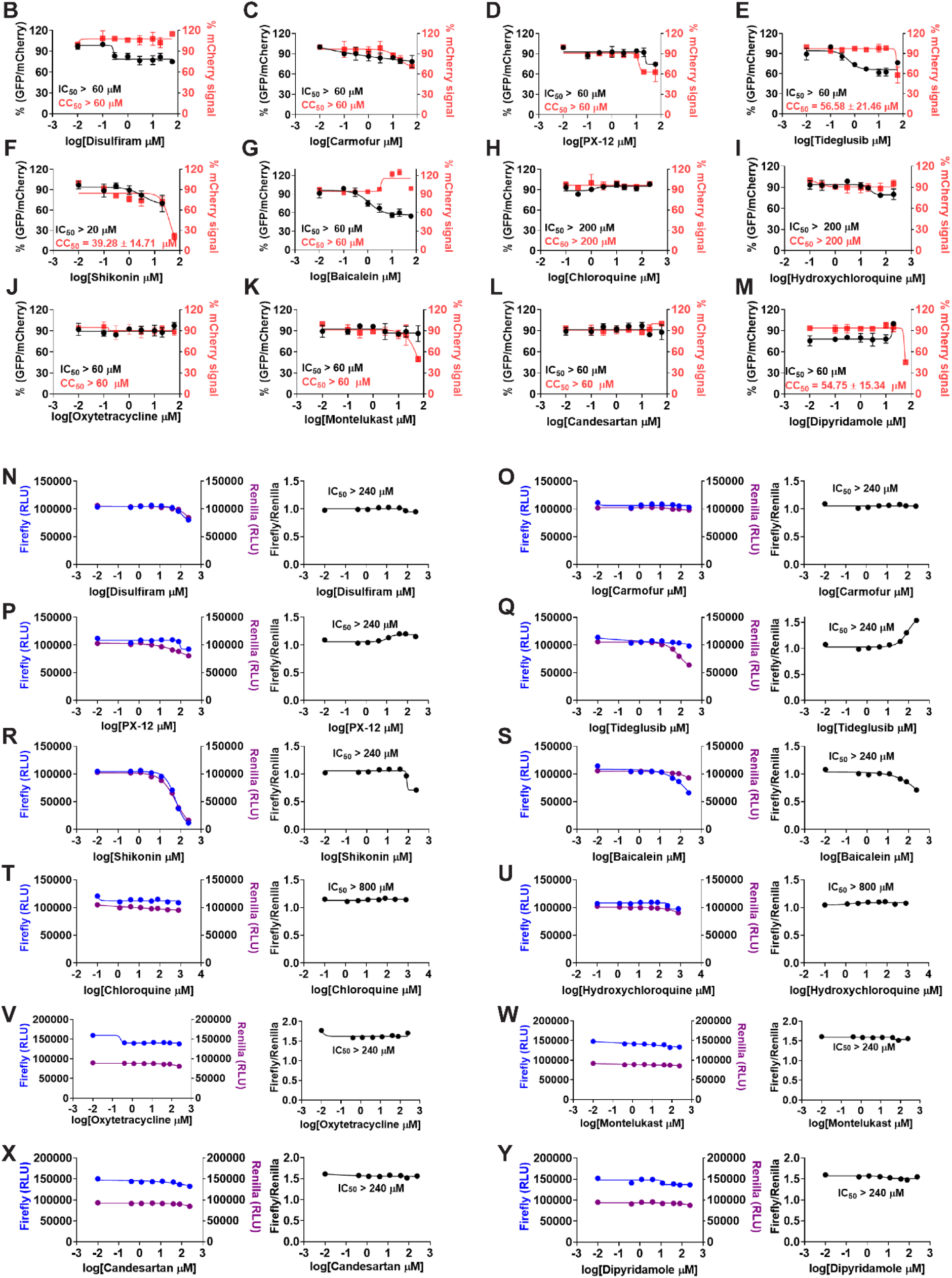
Validation/invalidation of disulfiram, carmofur, PX-12, tideglusib, shikonin, baicalein, chloroquine, hydroxychloroquine, oxytetracycline, montelukast, candesartan, and dipyridamole as SARS CoV-2 M^pro^ inhibitors using the Flip-GFP assay and Protease-Glo luciferase assay. (A) Representative images from the Flip-GFP-M^pro^ assay. (B-E) Dose−response curve of the ratio of GFP/mCherry fluorescent signal for disulfiram (B), carmofur (C), PX-12 (D), tideglusib (E), shikonin (F), baicalein (G), chloroquine (H), hydroxychloroquine (I), oxytetracycline (J), montelukast (K), candesartan (L), and dipyridamole (M); mCherry signal alone was used to normalize protein expression level or calculate compound cytotoxicity. (N-Y) Protease-Glo luciferase assay results of disulfiram (N), carmofur (O), PX-12 (P), tideglusib (Q), shikonin (R), baicalein (S), chloroquine (T), hydroxychloroquine (U), oxytetracycline (V), montelukast (W), candesartan (X), and dipyridamole (Y). Left column showed Firefly and Renilla luminescence signals in the presences of increasing concentrations of disulfiram, carmofur, PX-12, tideglusib, shikonin, baicalein, chloroquine, hydroxychloroquine, oxytetracycline, montelukast, candesartan, and dipyridamole; Right column showed dose−response curve plots of the ratio of FFluc/Rluc luminescence. Renilla luminescence signal alone was used to normalize protein expression level.

Similar to disulfiram, carmofur, PX-12 and tideglusib, which were previously claimed by Yang et al as M^pro^ inhibitors, showed no inhibitory activity in either the Flip-GFP or Protease-Glo luciferase assay (Fig. 5A, C, D, E, O, P, Q), which is consistent with their lack of inhibition in the FRET assay in the presence of DTT^35^.

Shikonin and baicalein are polyphenol natural products with known polypharmacology. Both compounds showed no inhibition in either the Flip-GFP or the Protease-Glo luciferase assay (Fig. 5A, F, G, R, S), suggesting they are not M^pro^ inhibitors. These two compounds were previously reported to inhibit SARS-CoV-2 M^pro^ in the FRET assay^8^ and had antiviral activity against SARS-CoV-2 in Vero E6 cells. However, our recent study showed that shikonin had no inhibition against SARS-CoV-2 M^pro^ in the FRET assay in the presence of DTT^35^. Studies from Deval et al using FRET assay and mass spectrometry assay reached the same conclusion. X-ray crystal structure of SARS-CoV-2 M^pro^ in complex with Shikonin showed that shikonin binds to the active site in a non-covalent manner.^9^

In addition to the proposed mechanism of action of M^pro^ inhibition, Schinazi et al showed that baicalein and baicalin inhibit the SARS-CoV-2 RNA-dependent RNA polymerase^62^. Overall, shikonin and baicalein are not M^pro^ inhibitors and the antiviral activity of baicalein against SARS-CoV-2 might involve other mechanisms.

A recent study from Luo et al identified several known drugs as SARS-CoV-2 M^pro^ inhibitors from a virtual screening^63^. The identified compounds include chloroquine (IC_50_ = 3.9 ± 0.2 μM; K_i_ = 0.56 ± 0.12 μM), hydroxychloroquine (IC_50_ = 2.9 ± 0.3 μM; K_i_ = 0.36 ± 0.21 μM), oxytetracycline (IC_50_ = 15.2 ± 0.9 μM; K_i_ = 0.99 ± 0.06 μM), montelukast (IC_50_ = 7.3 ± 0.5 μM; K_i_ = 0.48 ± 0.04 μM), candesartan (IC_50_ = 2.8 ± 0.3 μM; K_i_ = 0.18 ± 0.02 μM), and dipyridamole (K_i_ = 0.04 ± 0.001 μM). The discovery of chloroquine and hydroxychloroquine as M^pro^ inhibitor was particularly intriguing. Several high-throughput screenings have been conducted for M^pro24, 64^, and chloroquine and hydroxychloroquine were not among the list of active hits. In our follow up study, we found that none of the identified hits reported by Luo et al inhibited M^pro^ either with or without DTT in the FRET assay^30^. In corroborate with our previous finding, the Flip-GFP and Protease-Glo luciferase assays similarly confirmed the lack of inhibition of these compounds against M^pro^ (Fig. 5A, H-M, T-Y). Therefore, it can be concluded that chloroquine, hydroxychloroquine, oxytetracycline, montelukast, candesartan, and dipyridamole are not SARS-CoV-2 M^pro^ inhibitors. Other than the claims made by Luo et al, no other studies have independently confirmed these compounds as M^pro^ inhibitors.

## 3. CONCLUSION

The M^pro^ is perhaps the most extensive exploited drug target for SARS-CoV-2. A variety of drug discovery techniques have been applied to search for M^pro^ inhibitors. Researchers around the world are racing to share their findings with the scientific community to expedite the drug discovery process. However, the quality of science should not be compromised by the speed. The mechanism of action of drug candidates should be thoroughly characterized in biochemical, binding, and cellular assays. Pharmacological characterization should address both target specificity and cellular target engagement. For target specificity, the drug candidates can be counter screened against unrelated cysteine proteases such as the viral EV-A71 2A^pro^, EV-D68 2A^pro^, the host cathepsins B, L, and K, caspase, calpains I, II, and III, and etc. Compounds inhibit multiple cysteine proteases non-discriminately are most likely promiscuous compounds that act through redox cycling, inducing protein aggregation, or alkylating catalytic cysteine residue C145. For cellular target engagement, the Flip-GFP and Protease-Glo luciferase assays can be applied. Both assays are performed in the presence of competing host proteins at the cellular environment. Collectively, our study reaches the following conclusions: 1) for validated M^pro^ inhibitors, the IC_50_ values with and without reducing reagent should be about the same in the FRET assay; 2) validated M^pro^ inhibitors should show consistent results in the FRET assay, thermal shift binding assay, and the Protease-Glo luciferase assay. For compounds that are not cytotoxic, they should also be active in the Flip-GFP assay; 3) compounds that have antiviral activity but lack consistent results from the FRET, thermal shift, Flip-GFP, and Protease-Glo luciferase assays should not be classified as M^pro^ inhibitors; 4) compounds that non-specifically inhibit multiple unrelated viral or host cysteine proteases are most likely promiscuous inhibitors that should be triaged. 5) X-ray crystal structures cannot be used to justify the target specificity or cellular target engagement. Promiscuous compounds have been frequently co-crystallized with M^pro^ including ebselen, carmofur, and shikonin (Table 2).

Overall, we hope our studies will promote the awareness of the promiscuous SARS-CoV-2 M^pro^ inhibitors and call for more stringent hit validation.

## 4. MATHODS AND MATERIALS

### Protein Expression and Purification

The tag-free SARS CoV-2 M^pro^ protein with native N- and C-termini was expressed in pSUMO construct as described previously^3^.

### Enzymatic Assays

The FRET-based protease was performed as described previously^2^. Briefly, 100 nM of M^pro^ protein in the reaction buffer containing 20 mM HEPES, pH 6.5, 120 mM NaCl, 0.4 mM EDTA, 4 mM DTT, and 20% glycerol was incubated with serial concentrations of the testing compounds at 30 °C for 30 min. The proteolytic reactions were initiated by adding 10 μM of FRET-peptide substrate (Dabcyl-KTSAVLQ/SGFRKME(Edans)) and recorded in Cytation 5 imaging reader (Thermo Fisher Scientific) with 360/460 filter cube for 1 hr. The proteolytic reaction initial velocity in the presence or absence of testing compounds was determined by linear regression using the data points from the first 15 min of the kinetic progress curves. IC_50_ values was calculated by a 4-parameter dose−response function in prism 8.

### Thermal shift assay (TSA)

Direct binding of testing compounds to SARS CoV-2 M^pro^ protein was evaluated by differential scanning fluorimetry (DSF) using a Thermal Fisher QuantStudio 5 Real-Time PCR System as previously described^2^. Briefly, SARS CoV-2 M^pro^ protein was diluted into reaction buffer to a final concentration of 3 μM and incubated with 40 μM of testing compounds at 30 °C for 30 min. DMSO was included as a reference. SYPRO orange (1×, Thermal Fisher, catalog no. S6650) was added, and the fluorescence signal was recorded under a temperature gradient ranging from 20 to 95 °C with incremental step of 0.05 °C s^−1^. The melting temperature (T_m_) was calculated as the mid log of the transition phase from the native to the denatured protein using a Boltzmann model in Protein Thermal Shift Software v1.3. ΔT_m_ was the difference between T_m_ in the presence of testing compounds and T_m_ in the presence of DMSO.

### Flip-GFP M^pro^ Assay

The construction of FlipGFP-M^pro^ plasmid was described previously^11^. The assay was carried out as follows: 293T cells were seeded in 96-well black, clear bottomed Greiner plate (catalog no. 655090) and incubated overnight to reach 70− 90% confluency. 50 ng of FlipGFP-M^pro^ plasmid and 50 ng SARS CoV-2 M^pro^ expression plasmid pcDNA3.1 SARSCoV-2 M^pro^ were transfected into each well with transfection reagent TransIT-293 (Mirus catalog no. MIR 2700) according to the manufacturer’s protocol. Three hours after transfection, 1 μL of testing compound was directly added to each well without medium change. Two days after transfection, images were taken with Cytation 5 imaging reader (Biotek) using GFP and mCherry channels via 10× objective lens and were analyzed with Gen5 3.10 software (Biotek). The mCherry signal alone in the presence of testing compounds was utilized to evaluate the compound cytotoxicity.

### Protease-Glo luciferase assay

pGlosensor-30F DEVD vector was obtained from Promega (Catlog no. CS182101). pGloSensor-30F M^pro^ plasmid was generated by replacing the original caspase cutting sequence (DEVDG) was with SARS CoV-2 M^pro^ cutting sequence (AVLQ/SGFR) from BamHI/HindIII sites. The DNA duplex containing M^pro^ cutting sequence was generated by annealing two 5’-phosphoriated primers: forward: GATCCGCCGTGCTGCAGAGCGGCTTCAGA; and reverse: AGCTTCTGAAGCCGCTCTGCAGCACGGCG. Protease-Glo luciferase assay was carried out as follows: 293T cells in 10 cm culture dish were transfected with pGlosensor-30F M^pro^ plasmid in the presence of transfection reagent TransIT-293 (Mirus catalog no. MIR 2700) according to the manufacturer’s protocol. 24 hrs after transfection, cells were washed with PBS once, then each dish of cells was lysed with 5 ml of PBS+ 1% Trition-X100; cell debris was removed by centrifuge at 2000g for 10 min. Cell lysates was freshly frozen to -80 °C until ready to use. During the assay, 20 μl cell lysate was added to each well in 96-well flat bottom white plate (Fisherbrand Catalog no. 12566619), then 1 μl of testing compound or DMSO was added to each well and mixed at room temperature for 5 min. 5 μl of 200 nM *E. Coli* expressed SARS CoV-2 M^pro^ protein was added to each well to initiate the proteolytic reaction (the final M^pro^ protein concentration is around 40 nM). The reaction mix was further incubated at 30 °C for 30 min. The firefly and renilla luciferase activity were determined with Dual-Glo Luciferase Assay according to manufacturer’s protocol (Promega Catalog no. E2920). The efficacy of testing compounds against M^pro^ was evaluated by plotting the ratio of firefly luminescence signal over the renilla luminescence signal versus the testing compound concentrations with a 4-parameter dose−response function in prism 8.

### Antiviral assay in Calu-3 cells

The antiviral assay was performed as previously described^65^. Calu-3 cells (ATCC, HTB-55) were plated in 384 well plates and grown in Minimal Eagles Medium supplemented with 1% non-essential amino acids, 1% penicillin/streptomycin, and 10% FBS. The next day, 50 nL of compound in DMSO was added as an 8-pt dose response with three-fold dilutions between testing concentrations in triplicate, starting at 40 μM final concentration. The negative control (DMSO, n=32) and positive control (10 μM Remdesivir, n=32) were included on each assay plate. Calu-3 cells were pretreated with controls and testing compounds (in triplicate) for 2 hours prior to infection. In BSL-3 containment, SARS-CoV-2 (isolate USA-WA1/2020) diluted in serum free growth medium was added to plates to achieve an MOI of 0.5. Cells were incubated with compounds and SARS-CoV-2 virus for 48 hours. Cells were fixed and then immunostained with anti-dsRNA (J2) and nuclei were counterstained with Hoechst 33342 for automated microscopy. Automated image analysis quantifies the number of cells per well (toxicity) and the percentage of infected cells (dsRNA+ cells/cell number) per well. SARS-CoV-2 infection at each drug concentration was normalized to aggregated DMSO plate control wells and expressed as percentage-of-control (POC=% Infection _sample_/Avg % Infection _DMSO cont_). A non-linear regression curve fit analysis (GraphPad Prism 8) of POC Infection and cell viability versus the log_10_ transformed concentration values to calculate EC_50_ values for Infection and CC_50_ values for cell viability. Selectivity index (SI) was calculated as a ratio of drug’s CC_50_ and EC_50_ values (SI = CC_50_/IC_50_).

## Author contributions

Chunlong Ma performed the Flip-GFP assay, Protease-Glo luciferase assay, and thermal shift assay with the assistance from Haozhou Tan. Juliana Choza and Yuying Wang expressed the M^pro^ and performed the FRET assay. Jun Wang wrote the draft manuscript with the input from others; Jun Wang submitted this manuscript on behalf of other authors.

## Acknowledgments

This research was supported by the National Institute of Allergy and Infectious Diseasess of Health (NIH) (grants AI147325, AI157046, and AI158775) and the Arizona Biomedical Research Commission Centre Young Investigator grant (ADHS18-198859) to J. W. The SARS-CoV-2 antiviral assay in Calu-3 cells was conducted by Drs. David Schultz and Sara Cherry at the University of Pennsylvania through the NIAID preclinical service under a non-clinical evaluation agreement.

